# Bacteria isolated from bengal cat (*Felis catus* × *Prionailurus bengalensis*) anal sac secretions produce volatile compounds associated with animal signaling

**DOI:** 10.1101/625079

**Authors:** Mei S. Yamaguchi, Holly H. Ganz, Adrienne W. Cho, Thant H. Zaw, Guillaume Jospin, Mitchell M. McCartney, Cristina E. Davis, Jonathan A. Eisen, David A. Coil

**Author notes:** Corresponding author (JE).

## Abstract

Anal sacs are an important odor producing organ found across the mammalian Order Carnivora. Secretions from the anal sac may be used as chemical signals by animals for behaviors ranging from defense to species recognition to signaling reproductive status. In addition, a recent study suggests that domestic cats utilize short-chain free fatty acids in anal sac secretions for individual recognition. The fermentation hypothesis is the idea that symbiotic microorganisms living in association with animals contribute to odor profiles used in chemical communication and that variation in these chemical signals reflects variation in the microbial community. Here we examine the fermentation hypothesis by characterizing volatile organic compounds (VOC) and bacteria isolated from anal sac secretions collected from a male bengal cat, a cross between a domestic cat and wild leopard cat (*Felis catus* × *Prionailurus bengalensis*).

Both left and right anal sacs of a male bengal cat were manually expressed (emptied) and collected. Half of the material was used to culture bacteria or to extract bacterial DNA and other half was used for VOC analysis. DNA was extracted from the anal sac secretions and used for a 16S rRNA gene sequence based characterization of the microbial community. Additionally, some of the material was plated out in order to isolate bacterial colonies. The same three taxa, *Bacteroides fragilis*, *Tessaracoccus*, and *Finegoldia magna* were abundant in the 16S rRNA gene sequence data and also isolated by culturing. Using Solid Phase Microextraction (SPME) gas chromatography-mass spectrometry (GC-MS), we tentatively identified 52 compounds from bengal cat anal sac secretions and 67 compounds from cultures of the three bacterial isolates chosen for further analysis.. Among 67 compounds tentatively identified from bacteria isolates, 52 were also found in the anal sac secretion.

We show that the bacterial community in the anal sac consists primarily of only a few abundant taxa and that isolates of these taxa produce numerous volatiles that are found in the combined anal sac volatile profile. Many of these volatiles are found in anal sac secretions from other carnivorans, and are also associated with known bacterial biosynthesis pathways. This supports the fermentation hypothesis and the idea that the anal sac is maintained at least in part to house bacteria that produce volatiles for the host.

## Introduction

Anal sacs are an odor producing organ common to many mammals, including members of the Order Carnivora (carnivorans) [1,2]. In carnivorans, anal sacs are two small structures found on each side of the anus [3], located between the internal and external sphincter muscles. The interior walls of each sac are lined with sebaceous and apocrine glands, and the anal sac secretes a foul smelling, oily substance that ranges in color from yellow to brown [4]. Scent secretions and marking behaviors play critical roles in animal communication. Anal sac secretions are used for defense by the hooded skunk (*Mephitis macroura*) [5] and the honey badger (*Mellivora capensis*) [6], territory marking by the spotted hyena (*Crocuta crocuta*) [7] and the wolf (*Canis lupus*) [8,9], individual identification by the ferret (*Mustela furo*) [10,11], the mongoose (*Herpestes auropunctatus*) [12], the giant panda (*Ailuropoda melanoleuca*) [13], striped hyena (*Hyaena hyaena*) [14] and the spotted hyena (*C. crocuta*) [13,15] and sex recognition by the brown bear (*Ursus arctos*) [16], the giant panda (*A. melanoleuca*) [17], and some mustelids (*Mustela* spp.) [11,18]. In domestic cats (*Felis catus*) and meerkats (*Suricata suricatta*) [19], anal sac secretions are used for territorial marking, and such secretions may have information about sex, reproductive state, and recognition of individuals [1,20]. Further, these chemical signals are species specific and chemical signals from scent glands in the Felidae were found to retain a phylogenetic signal [21].

The chemical composition of anal sac secretions has been analyzed in a number of animals in the Carnivora. Studies in the cheetah (*Acinonyx jubatus*), red fox (*Vulpes vulpes*), dog (*Canis familiaris*), coyote (*Canis latrans*), wolf (*C. lupus*), lion (*Panthera leo*), and mongoose (*H. auropunctatus*) have identified volatile short-chain free fatty acids, such as acetic acid, propanoic acid, and butanoic acid as being partially responsible for the odors [22–28]. The nature of these constituents led to the suggestion that they may be metabolites produced by bacteria in the sac from available substrates [22].

The fermentation hypothesis posits that bacteria metabolize secretions and produce volatile organic compounds, such as hydrocarbons, fatty acids, wax esters, and sulfur compounds [15,16,29] that are used in communication by the host [30,31]. Evidence in support of this hypothesis links bacterial action to specific, olfactory-mediated host behavior or to the production of certain odorants. For example, researchers have shown that trimethylamine, an odorant that plays a key role in mouse (*Mus musculus*) reproduction, requires commensal bacteria for its production [24]. Researchers have also inhibited odorant production in Indian mongooses (*H. auropunctatus*) and European hoopoes (*Upupa epops*) by treating the animals’ scent glands with antibiotics [30,32].

In this study, we investigated the fermentation hypothesis by focusing in detail on bacterial isolates collected from a single animal, available at the time of our study. We studied a bengal cat, a hybrid between the Asian leopard cat (*P. bengalensis*) and the domestic cat (*F. catus*). We collected anal sac secretions in order to characterize their chemical profile and analyze bacterial community composition. Then we isolated and identified those bacteria that could be cultivated under anaerobic conditions from these samples. Volatiles produced by these isolates were identified and compared to those found in the anal sac secretions. This is the first study in felines to demonstrate that bacteria isolated from anal sacs produce key volatile compounds found in anal sac secretions.

## Materials and methods

### Sample collection

With the owner’s consent, both left and right anal sacs were manually expressed in a male bengal cat (*F. catus* × *P. bengalensis*) by a veterinarian at the Berkeley Dog and Cat Hospital in Berkeley, CA. Samples of anal sac secretions were collected using Puritan cotton swabs and placed in 2 mL screw cap tubes. Three swabs (2 samples and one blank control) were used for 16S rRNA gene sequencing (placed into 100% ethanol), three swabs were used for GC/MS (plus two media controls and two jar blanks), and two swabs were used for culturing.

### DNA extraction and 16S rRNA gene sequencing and analysis

Swabs of the anal sac secretions were placed in 100% ethanol prior to extraction. Genomic DNA was extracted using the MoBio PowerSoil DNA Isolation kit (MoBio, Carlsbad, CA, USA). Samples were transferred to bead tubes containing C1 solution and incubated at 65 °C for 10 min, followed by 3 min of bead beating. The remaining extraction protocol was performed as directed by the manufacturer.

DNA samples were sent to the Integrated Microbiome Resource, Centre for Comparative Genomics and Evolutionary Bioinformatics, Dalhousie University for sequencing. Bacterial diversity was characterized via amplification by a PCR enrichment of the 16S rRNA gene (V4-V5 region) using primers 515F and 926R (Walters et al. 2016). After an initial denaturation step at 98 °C for 30 sec, we ran 30 cycles of the following PCR protocol: 98 °C for 10 s, 55 °C for 30 s and 72 °C for 30 s, followed by final extension at 72 °C for 4 m 30 s, and a final hold at 4 °C [33]. Prior to sequencing, the amount of input DNA per sample was normalized using a SequalPrep Normalization Plate, following the standard protocol (ThermoFisher Scientific, Wilmington, DE, USA). The final library pool was quantified using the Qubit dsDNA HS assay (Invitrogen, Carlsbad, CA). We generated paired-end 300 bp sequencing reads of 16S PCR amplicons with multiple barcodes on the Illumina MiSeq, resulting in 450 bp amplicons of the V4 region using v3 chemistry. The samples were diluted to the appropriate concentration, spiked with 5% PhiX control library, and sequenced using an Illumina MiSeq instrument with the manufacturer’s standard 150 nucleotides paired-end dual-index sequencing protocol.

Raw, demultiplexed amplicon reads were processed using DADA2 v1.8, following the standard online tutorial [34]. The reads were trimmed down to 250 base pairs to remove low quality nucleotides. In addition, the quality of reads were ensured by trimming bases that did not satisfy a Q2 quality score. Reads containing Ns were discarded and we used two expected errors to filter the overall quality of the read (rather than averaging quality scores) [35]. Chimeric reads were also removed using DADA2 on a per sample basis. The remaining pairs of reads were merged into amplicon fragments and unique Amplicon Sequence Variants (ASVs) were identified. Reads that did not merge successfully were discarded. Upon completion of the DADA2 pipeline, all ASVs (n=52) that were found in the negative control swabs were removed from the analysis, only three of these ASVs were also found in the anal sac samples. ASVs were assigned taxonomy using the dada2 function “assignTaxonomy” and the Silva (NR v132) database [36–38]. One ASV that was assigned to “Eukaryotes” was removed. All ASVs with the same taxonomy (at the genus level) were grouped and then ranked by number of reads. No ASVs were assigned to mitochondria or chloroplast.

### Bacterial culturing and identification

Anal sac secretions were vortexed with 1 mL Phosphate Buffer Saline (PBS). Two serial 1:10 dilutions were performed and 100 µL of each dilution was plated onto Columbia Blood Agar (CBA) and Brain Heart Infusion (BHI). Plates were incubated anaerobically in a BD GasPak EZ Container System with packets of CO_2_ generator for 5 days at 37 °C. Morphologically distinct colonies were streaked for isolation on both CBA and BHI. The 16S rRNA gene was sequenced using Sanger sequencing using the 27F and 1391R primers. Taxonomy was assigned by the result of BLAST queries to the nr database at NCBI (excluding unnamed/environmental sequences), a species name was given in cases where the identity was >98% to only a single species.

### Bacterial volatile analysis

To extract volatiles from *Bacteroides fragilis* UCD-AAL1 and *Tessaracoccus sp.* UCD-MLA, cultures were grown in 5 mL BHI anaerobically for 24 hours at 37 °C. 100 µL of the culture was transferred into each of three Restek (Bellefonte, PA) tubes filled with 5 mL of BHI. Two jar blanks (no media or bacteria) and two BHI media-only blanks were used as controls. The same procedure was followed for *Finegoldia magna* UCD-MLG, except that cultures were grown and incubated in BHI supplemented with 5% defibrinated sheep blood (BBHI) anaerobically for 24 hours at 37°C.

Headspace extraction was performed with Solid Phase Microextraction (SPME) fibers (Part 57912-U, Sigma Aldrich) which had 50/30 μm thickness and DVB/CAR/PDMA coating. Two SPMEs were inserted into the headspace of each Restek tube prior to anaerobic incubation at 37 °C for 24 hours. SPME fibers were introduced by piercing the fibers through the septa insert of the lids and making sure that the fibers were injected but not touching the media containing the bacteria. An internal standard was introduced before sampling using 1 μL of the standard solution (10 mL/L of decane-d22 in ethanol) per jar.

For the anal sac samples, the swabs containing the anal sac fluid were placed in septa screw cap jars that each contained a SPME. After a 24 hour incubation period, the SPMEs were removed. Then we performed a liquid extraction of volatiles by adding 20 mL of methanol to the jars and incubating for 24 hours.

### GC-MS analysis

Chromatography occurred on a 7890 GC (Agilent Technologies Inc., Santa Clara, CA) with a ZB-WAX 30 m × 250 μm capillary column, coated with a 0.25 µm film stationary phase (Part 7HG-G007-11, 100% polyethylene glycol from Phenomenex, Torrance, CA) equivalent to DB-Wax or Carbowax. Helium was used as the carrier gas at 1 ml/min in constant flow mode. The inlet was set to 260 ℃ and SPMEs were splitlessly desorbed during the run. The oven temperature was programmed to increase from 40 ℃ (held for 5 min) to 110 °C at a rate of 5 °C min-1, and raise to 250 °C (held for 10 min) at a rate of 40 °C min−1. A transfer line set at 250 °C led to a 5977A mass spectrometer (Agilent Technologies Inc., Santa Clara, CA) with a solvent delay of 5 min. The MS swept from 50 to 500 m/z. The mass spectrometer was operated in the selected scan mode. The MS source was set to 230 °C and the MS quad set to 150 °C. A standard mix of C_8_-C_20_ alkanes was analyzed to calculate the Kovats Retention Indices and to monitor control of the instruments.

Methanolic extract of cat anal secretion was analyzed by GC-MS as tert-butyldimethylsilyl (TBDMS) derivatives (Knapp, 1979). 2 mL out of 20 mL was used in the analysis. Samples were placed in glass conical vials and dried, reacted with a mixture of 50 μL N-methyl-N-tert-butyldimethylsilyltrifluoroacetamide (MTBSTFA; Sigma-Aldrich Co. LLC., St Louis, MO, U.S.A.) and 50 μL acetonitrile at 60°C for an hour.

### GC/MS data analysis workflow

MassHunter Profinder B.08.00 (Agilent Technologies Inc.) was used to deconvolute, integrate and align the data. Peaks with amplitudes of less than 1000 counts were ignored. Compounds must have been present in at least 60% of replicates from one treatment to be included in statistical analyses. Statistical analysis was performed using GeneSpring MPP (Agilent Technologies, Inc.), where p ≤ 0.05 was used throughout to test for statistical significance using a T-test with Bonferroni correction. Tentative compound identification was based on the combined comparing mass spectra to the NIST 2014 Library and by a comparison of the calculated matching of standard alkane retention indices (LRI) values, when available.

## Results and discussion

### 16S rRNA gene sequencing and bacteria identification

The 16S rRNA gene PCR sequencing and analysis of the feline anal sac showed that 98% of the reads that were placed into ASVs were assigned to six genera (Table 1). These ASVs generally represent genera that contain anaerobic members known to be associated with mammals. *Tessaracoccus* species are have been isolated in sediment and have also been found in the gut of mammals including rhinoceros and humans [39–43]. *Bacteroides* is a genus of bacteria also often associated with mammals [44,45]. *Anaerococcus*, *Peptoniphilus* and *Finegoldia* are all Gram Positive Anaerobic Cocci (GPAC) formerly part of the *Peptostreptophilus* genus and are found in mammalian guts and urinary tracts [46–48]. *Peptostreptococcus* is another mammalian-associated GPAC, with around 15 species in the genus. At least one member of the group has been found as an obligate anaerobic bacteria in cats and dogs [49,50].

**Table 1.** 16S rRNA gene based survey of the bengal cat anal sacs.

| Right Anal Sac |  | Left Anal Sac |  |  |
| --- | --- | --- | --- | --- |
| # of reads | % | # of reads | % | Genus |
| 15250 | 83% | 17840 | 83% | *Tessaracoccus* |
| 1532 | 8% | 1986 | 9% | *Anaerococcus* |
| 709 | 4% | 746 | 3% | *Finegoldia* |
| 325 | 2% | 390 | 2% | *Bacteroides* |
| 182 | 1% | 114 | 1% | *Peptoniphilus* |
| 117 | 1% | 111 | 1% | *Peptostreptococcus* |
| 18115 | 98% | 21187 | 98% | n/a |

Table shows the number of reads mapping to ASVs, summed by taxonomy at the genus level (e.g. everything with the same genus was collapsed into a single column). Not shown are any reads that were placed into ASVs for which the genera of the taxonomic assignments summed to less than 1% of the total number of sequencing reads.

We cultured 25 bacterial isolates from this anal sac and found only *Tessaracoccus*, *Escherichia*, *Bacteroides*, *Finegoldia*, and *Clostridium* isolates under the conditions used (TABLE 2). Of the cultured bacteria, three isolates were members of genera found in high abundance in the anal sac: *Tessaracoccus*, *Bacteroides fragilis*, and *Finegoldia magna*. We therefore chose to focus our experimental efforts on these genera.

**Table 2.** Identified isolates in this study

| Strain ID | Genus | Species |
| --- | --- | --- |
| UCD-MLI | *Bacteroides* | *fragilis* |
| UCD-AAL1 | *Bacteroides* | *fragilis* |
| UCD-MBD1A | *Clostridium* | *perfringens* |
| UCD-MBD2A | *Clostridium* | *perfringens* |
| UCD-MD1A | *Clostridium* | *perfringens* |
| UCD-MD1C | *Clostridium* | *perfringens* |
| UCD-MD1E | *Clostridium* | *perfringens* |
| UCD-MD2A | *Clostridium* | *perfringens* |
| UCD-MLB | *Escherichia* | *coli* |
| **UCD-MLG** | ***Finegoldia*** | ***magna*** |
| **UCD-MLA** | ***Tessaracoccus*** | **sp.** |
| UCD-MLH | *Tessaracoccus* | sp. |
| UCD-ACL3 | *Tessaracoccus* | sp. |
| UCD-AD1A | *Tessaracoccus* | sp. |
| UCD-AD2B1 | *Tessaracoccus* | sp. |
| UCD-AD2B2 | *Tessaracoccus* | sp. |
| UCD-AD2B3 | *Tessaracoccus* | sp. |
| UCD-AD3C | *Tessaracoccus* | sp. |
| UCD-MLE | *Tessaracoccus* | sp. |
| UCD-MLF | *Tessaracoccus* | sp. |
| UCD-MBD1B | *Tessaracoccus* | sp. |
| UCD-MBD1C | *Tessaracoccus* | sp. |
| UCD-MBD1D | *Tessaracoccus* | sp |
| UCD-MBD1E | *Tessaracoccus* | sp |
| UCD-MBD3A | *Tessaracoccus* | sp |
| UCD-MD3B | *Tessaracoccus* | sp |

All identified isolates from the anal sacs used in this study, bolded taxa are those for which we generated volatile data. Identifications were made from BLAST results of Sanger sequencing of the 16S rRNA gene.

**Table 3.**
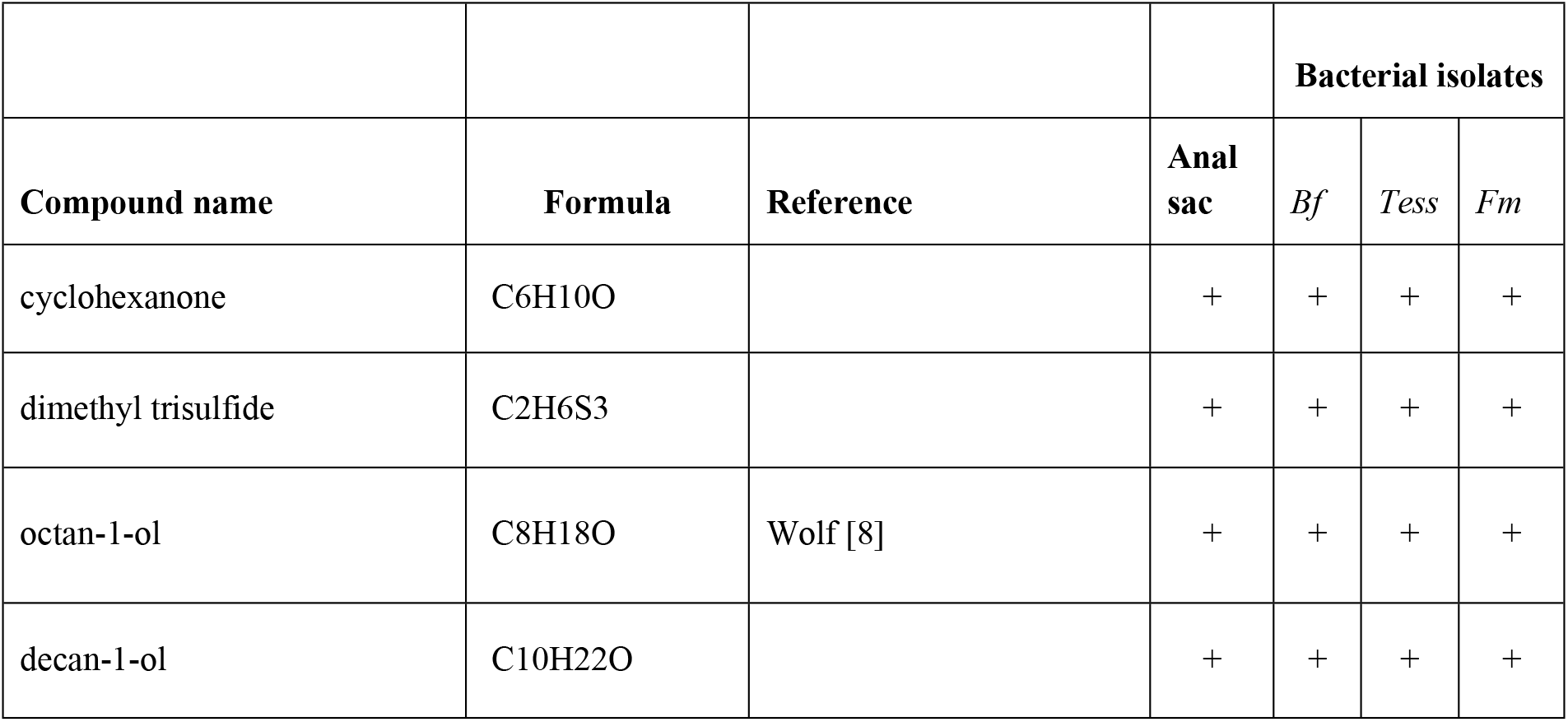
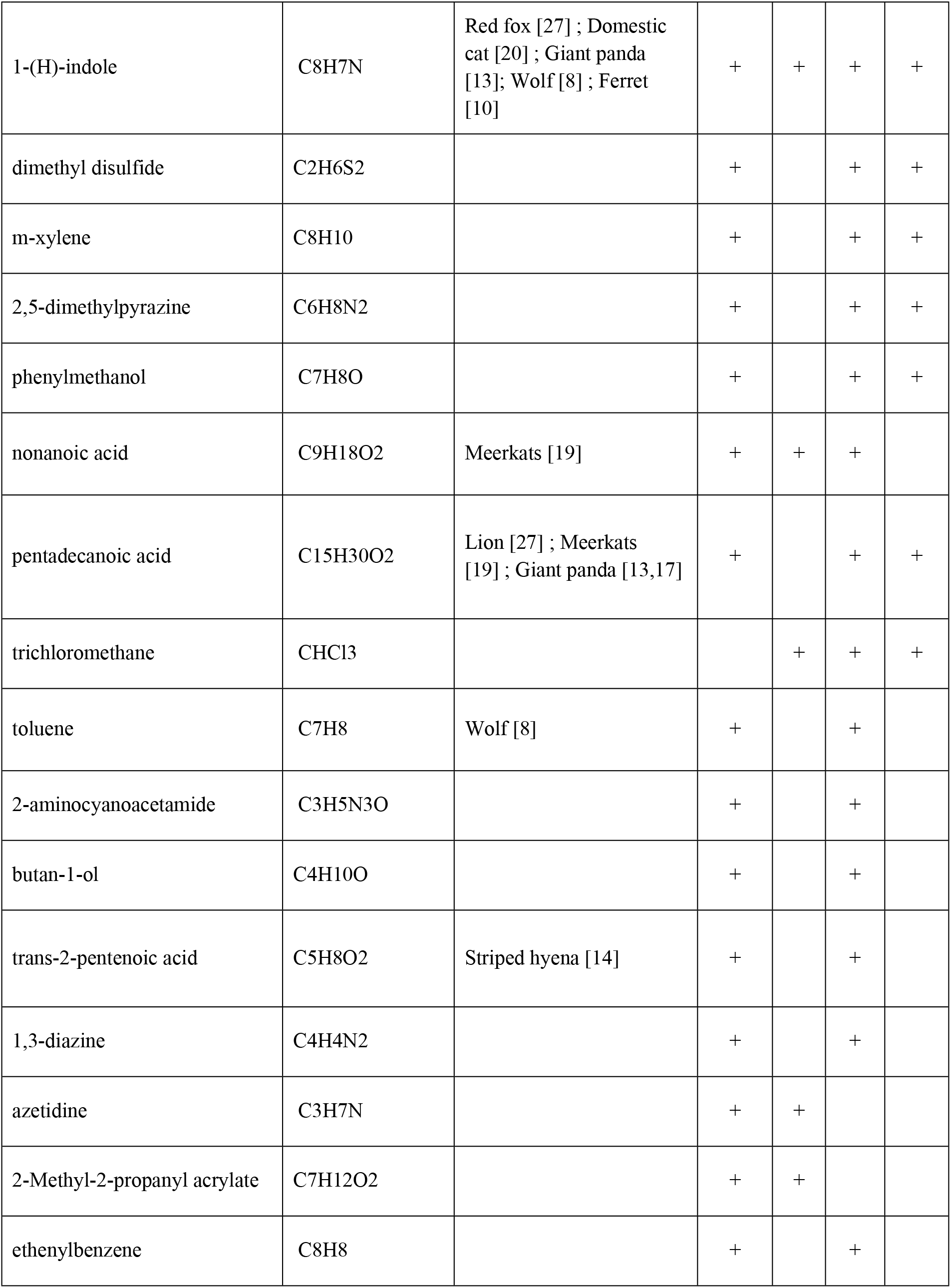
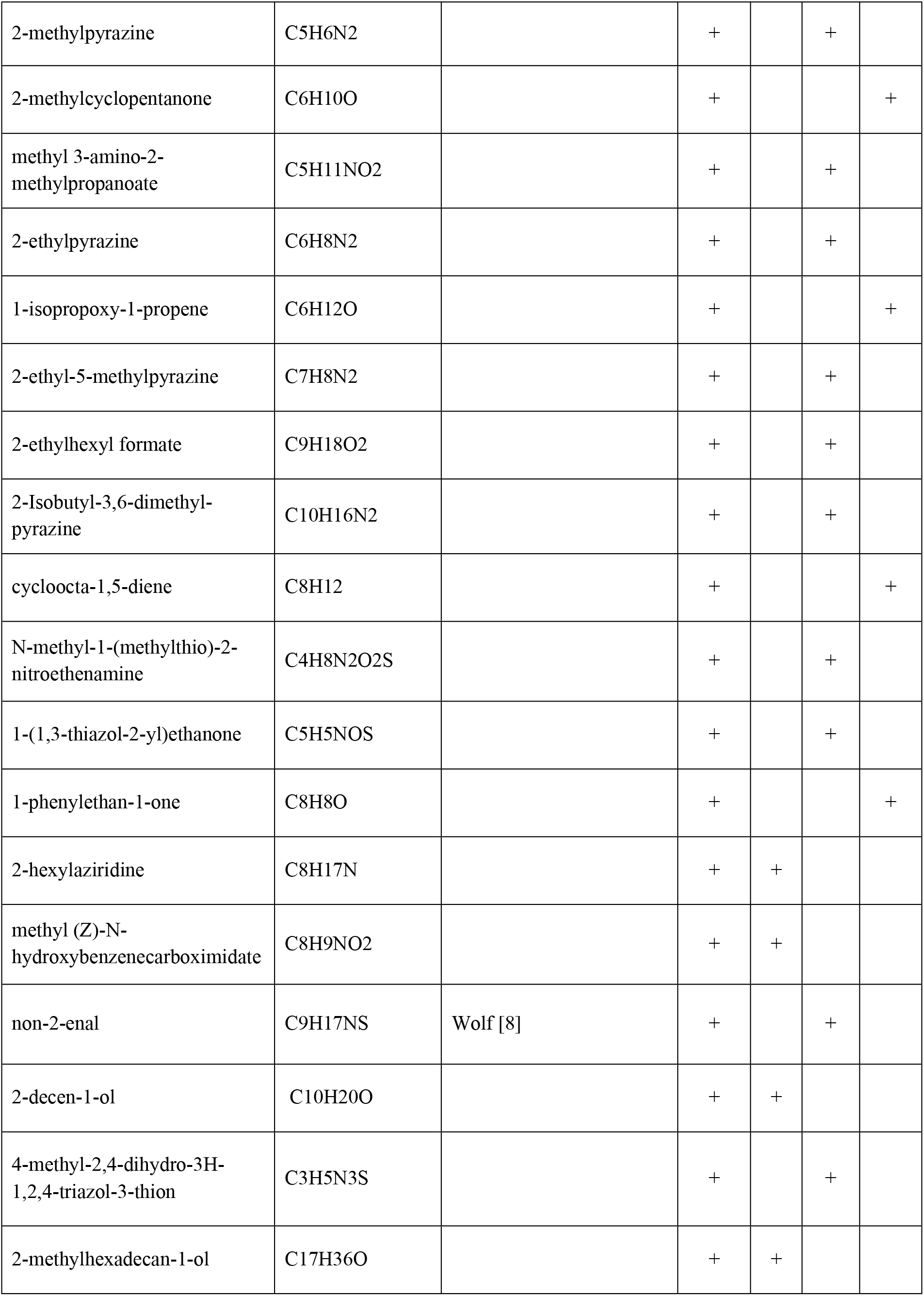
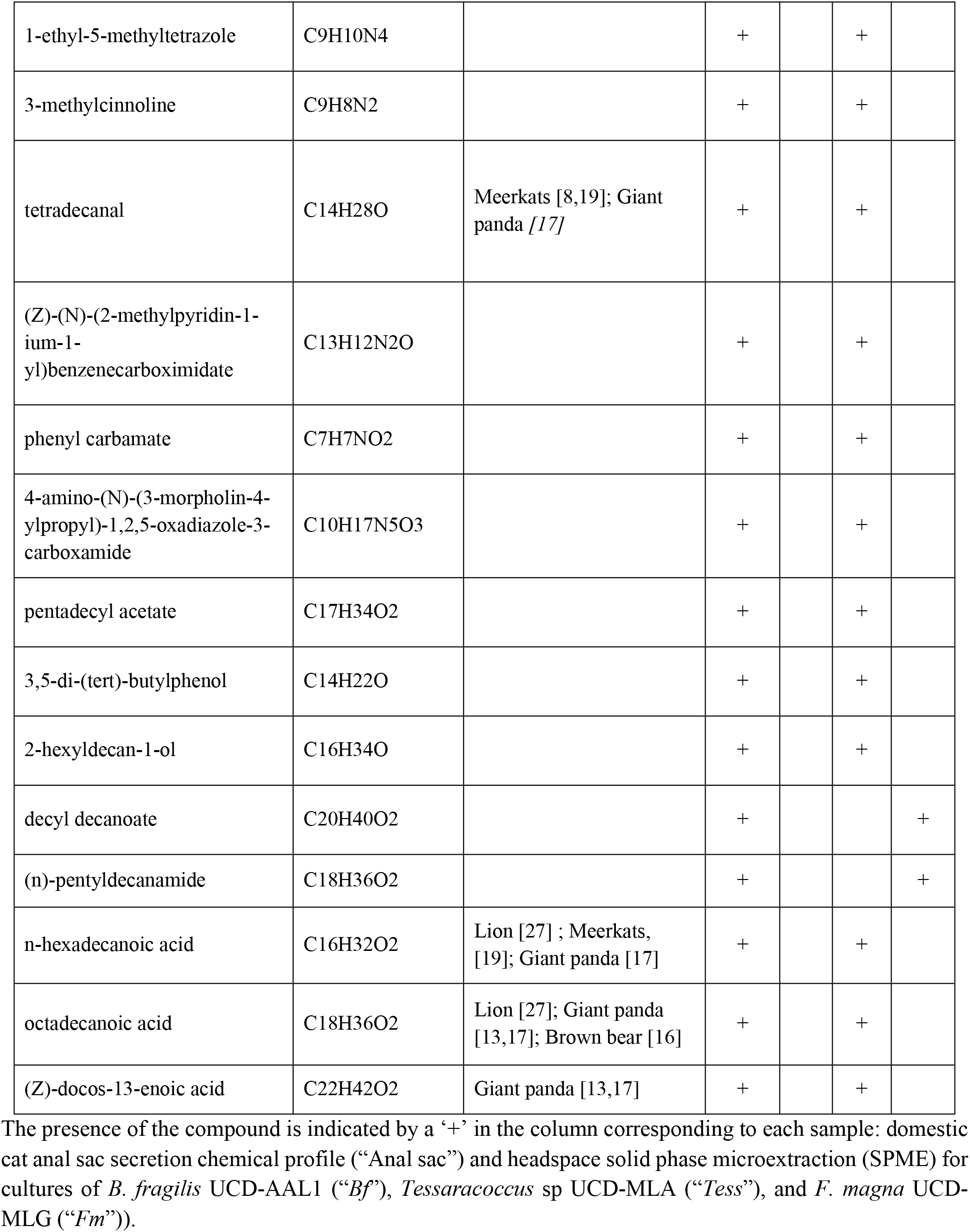
Tentatively identified compounds (compound name and formula listed) from cultures and anal sac.

*Tessaracoccus* is a genus in the Propionibacteriaceae family of the phylum Actinobacteria. The characterized species in this genus are facultatively anaerobic and Gram positive bacteria. Representatives from this genus have been isolated from a wide variety of environments, including rhinoceros feces [42], the gut of humans [41], metalworking fluid [51], marine sediment [39], and activated sludge biomass [52]. For our work on *Tessaracoccus*, we focused our efforts on a single isolate, *Tessaracoccus* sp. UCD-MLA. This full-length 16S rRNA gene sequence was 97% identical to multiple members of the genus and we could not assign a species ID with confidence [53].

*Bacteroides fragilis* is the type species of the genus *Bacteroides* in the Bacteroidaceae family (phylum Bacteroidetes) and is an obligate anaerobe [54]. It has previously been found in numerous places, including the oral cavity of cats [55], as well as the gut microbiome of dogs [56]. The 16S rRNA gene of our chosen isolate, *Bacteroides fragilis* UCD-AAL1, was 99% identical to representatives of this species at NCBI.

The only species in the genus of *Finegoldia* (Peptoniphilaceae family within the Firmicutes phylum), *Finegoldia magna*, is an obligately anaerobic Gram positive coccus that is part of the normal flora of the human gastrointestinal and genitourinary tracts [47] and has been previously found in cats [57] as well as dogs [58]. The 16S rRNA gene from our single isolate, *Finegoldia magna* UCD-MLG, was 99% identical to representatives of this species at NCBI.

### Anal sac secretion constituents and Bacteria isolates headspace SPME

We tentatively identified 127 compounds from the domestic cat anal gland secretion. Out of 127 tentatively identified compounds, 89 compounds were found in liquid extraction of anal secretion after TBDMS derivatization (S1 Table), and 52 compounds were measured by SPME-GC-MS in the anal sac secretion. These compounds were tentatively identified on the basis of the precise interpretation of its accurate mass spectra, MS fragmentation, and Kovats index information. These VOC metabolites were identified in the following compound chemical classes: heterocyclic compounds (12 %), alcohols (16 %), fatty acids (17 %), ketones (11 %), aromatic carbons (13 %), amines (9 %), aldehydes (7 %), esters (6 %).

A total of 67 compounds were tentatively identified from the SPME analysis of the three bacterial isolates (19 compounds from *B. fragilis* UCD-AAL1, 44 compounds from *Tessaracoccus* sp UCD-MLA, and 23 compounds from *F. magna* UCD-MLG) (Table 2). Among these 67 compounds, 52 compounds were also found in the anal sac secretion. 11 compounds (octan-1-ol, 1-(H)-indole, nonanoic acid, pentadecanoic acid, toluene, trans-2-pentenoic acid, non-2-enal, tetradecanal, n-hexadecanoic acid, octadecanoic acid, and (Z)-docos-13-enoic acid) found in the anal sac secretion have also been reported in other mammalian anal sac secretions [10,13,14,17,19,20,26,27]. Octan-1-ol and 1-(H)-indole were found in the anal sac secretion and in all three bacterial isolates. Octan-1-ol is a compound that produces a pungent odor and was previously reported in *C. lupus* anal sac secretion [26]. 1-(H)-indole is known to be widely distributed in the natural environment and can be produced by a variety of bacteria that have a strong fecal odor. 1-(H)-indole has also been found in various mammalian anal secretions, such as *V. vulpes [59]*, *F. catus [20]*, *A. melanoleuca [13]*, *C. lupus [26]*, and *M. furo [10]*. Toluene is an aromatic compound found naturally in petroleum and coal, and is a major component of gasoline [60]. Raymer *et. al.* reported toluene found in *C. lupus [26]*. Although anaerobic bacterial toluene degradation is a well known pathway [61], toluene biosynthesis is less common in bacteria. It is possible that our tentatively identified result could be a different compound but is likely to contain the alkyl benzene structure. Non-2-enal, and tetradecanal are chain aldehyde compounds. Non-2-enal was previously found in *C. lupus [26]* and tetradecanal was found in both *S. suricatta [19,27]*, and *A. melanoleuca [17]* anal sac secretions. It is likely these aldehydes are formed by bacterial oxidation of a ubiquitous fatty acid such as oleic acid [62]. Out of 52 compounds, six fatty acids were tentatively identified from our anal sac secretion; nonanoic acid, pentadecanoic acid, trans-2-pentenoic acid, n-hexadecanoic acid, octadecanoic acid, and (Z)-docos-13-enoic acid. Fatty acids are one of the most common compound groups known to be present in various animal anal sac secretions [13,14,16,17,19,27] and all of the fatty acids found in our anal sac secretion have been previously reported in other mammalian anal sac secretions. Nonanoic acid is a nine carbon fatty acid known to have unpleasant rancid odor, and previously found in *S. suricatta* anal sac secretions [19]. Pentadecanoic acid was found in *P. leo [27]*, *S. suricatta [19]*, and *A. melanoleuca [13,17]* anal sac secretions. Trans-2-pentenoic acid was found in *H. hyaena [14]* anal sac secretions. N-hexadecanoic acid was found in *P. leo [27]*, *S. suricatta [19]*, *A. melanoleuca [17]* anal sac secretions. (Z)-docos-13-enoic acid was found in *A. melanoleuca [13,17]* anal sac secretions. Fatty acids are a very common group biosynthesized by variety of organisms. The biogenesis typically starts with acetyl CoA, which is extended with malonate units to ultimately assemble fatty acids containing an even number of carbons [62]. All six fatty acids found in our anal sac secretion were also found in the *Tessaracoccus* sp UCD-MLA culture. It is possible that the highly abundant (in our sample) *Tessaracoccus* is largely contributing in the production of fatty acid compounds in this anal sac.

41 compounds found in our bengal cat anal sac secretion have not been previously described in an anal sac secretion. Cyclohexanone and dimethyl trisulfide were found from the anal sac secretion and also from all three bacteria isolates. Cyclohexanone is known to be generated by the cyclohexanol degradation metabolomic pathway, which is widely used by bacteria [63]. Including dimethyl trisulfide, volatile sulfur compounds (hydrogen sulfide, methanethiol, dimethyl sulfide, dimethyl disulfide) such methionine derived volatiles are often generated by bacteria [62] and have also been reported to be produced by the *Bacteroides* family [64]. Hydrocarbons, aliphatic alcohols, and ketones are volatiles most likely formed by modification of products of the fatty acid biosynthetic pathway [62]. Alkanes and methyl ketones are often known to be produced from decarboxylation in bacteria [62]. Methyl ketone group compounds (2-methyl cyclopentanone, 1-(1,3-thiazol-2-yl)ethanone, 1-phenylethan-1-one) were tentatively identified from the cat anal sac secretion and at least one of the bacteria isolates. Another final modification process of the fatty acid metabolomic pathway is the reduction of carboxy groups to aldehydes and into aliphatic alcohols. Several aliphatic alcohol (decan-1-ol, butan-1-ol, 2-methylhexadecan-1-ol, 2-hexyldecan-1-ol) were tentatively identified from the anal sac secretion and from bacteria isolates. Butan-1-ol and its bishomologues up to C16 have been found in different combinations in several bacteria [62,65,66]. Aromatic compounds are common natural products in plants but also known to be produced by bacteria [62]. Especially among the aromatic alcohols, such phenols, 2-penylethanol is one of the most aromatic compound produced by diverse bacteria. In our present study, aromatic alcohols, such as phenyl methanol, and 3,5-di-(tert)-buthylphenol were tentatively identified from both anal sac and bacterial isolates. These aromatic alcohols can be assumed to be generated by the shikimate pathway which is present only in microorganisms and plants, never in animals [67].

## Conclusion

To our knowledge, this is the first study examining either the VOC profile of domestic feline anal sacs or the VOC profiles of associated bacteria. We show that these particular feline anal sacs are dominated by only a few taxa, most of which are easily culturable under anaerobic conditions. These bacteria produce the majority of the identified volatiles in the total anal sac scent profile. Our preliminary identification of these volatiles is supported by the existence of known bacterial metabolic pathways for the majority of these compounds. Together these results support the fermentation hypothesis and suggest that further characterization of the anal sac microbial community, as well as the VOC’s produced therein could potentially shed light on the potentially symbiotic relationship between these microbes and their host.

## Acknowledgments

We thank Leah K. Isaacson, DVM for expressing the anal sacs and providing these samples to us. The authors would also like to thank Petra Dahms, Dana De Vries, and Alex Martin who worked on the initial stages of this project. We thank Alexandria Falcon for contributing her technical expertise.

Partial funding for this project was provided by various sources, including: the KittyBiome citizen science project, NIH training grant award T32 HL007013 [MSY], NIH award U01 EB0220003-01 (CED); NIH National Centre for Advancing Translational Sciences (NCATS) through award UL1 TR000002 (CED); NIH award UG3-OD023365 (CED); and NIH award 1P30ES023513-01A1 (CED). The contents of this manuscript are solely the responsibility of the authors and do not necessarily represent the official views of the funding agencies.

## Animal Use and Care

No laboratory animals were used in this research. The owner of a male bengal cat volunteered to have her cat’s anal sacs manually expressed by a veterinarian at the Berkeley Dog and Cat Hospital in Berkeley, CA.

## Availability of Supporting Data

All raw sequencing data has been deposited at NCBI under BioProject PRJNA533859. All data analysis, supporting files and intermediate analysis files are available at DASH (doi:10.25338/B8JG7X).

## Supporting information

**S1 Table.**
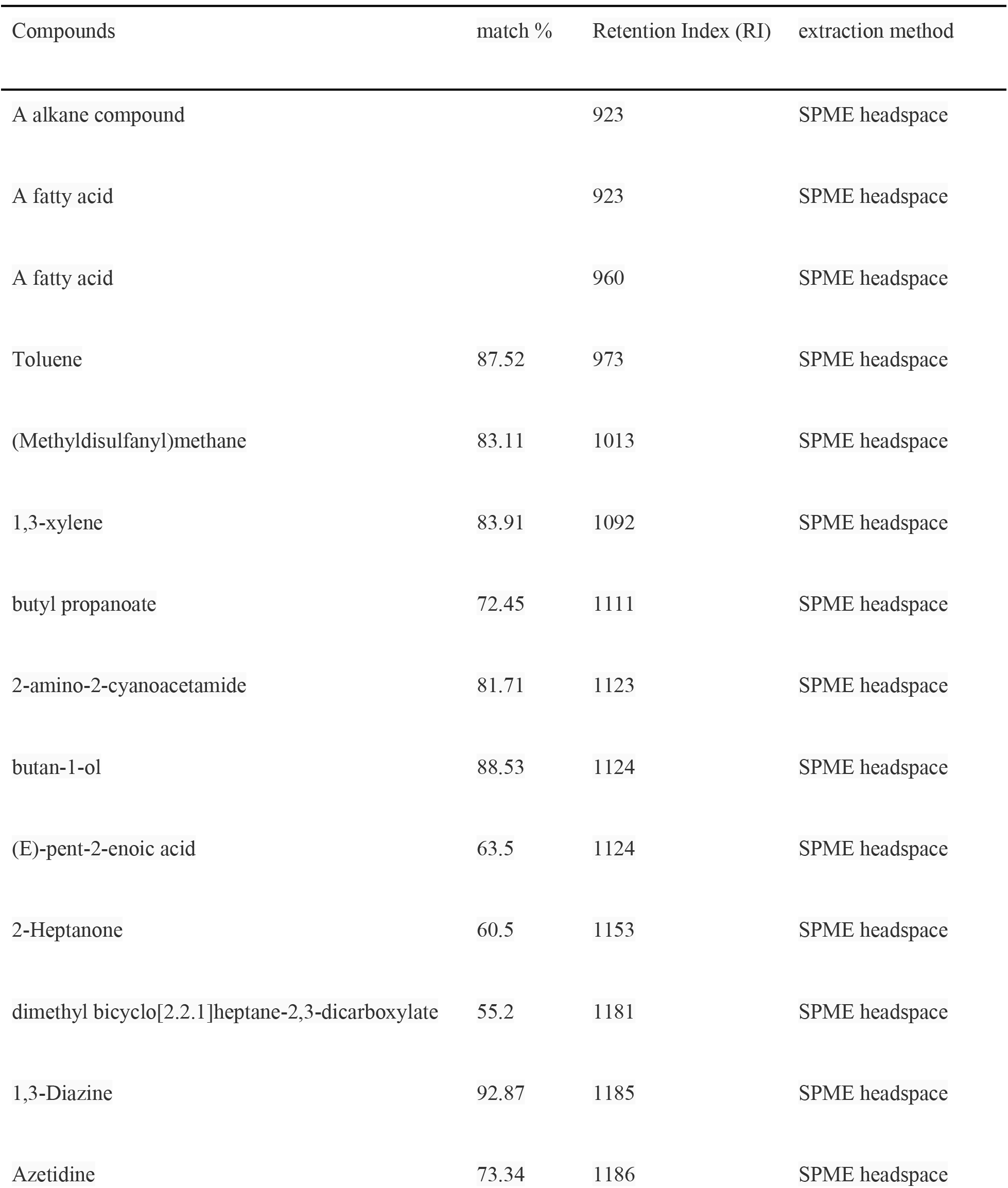
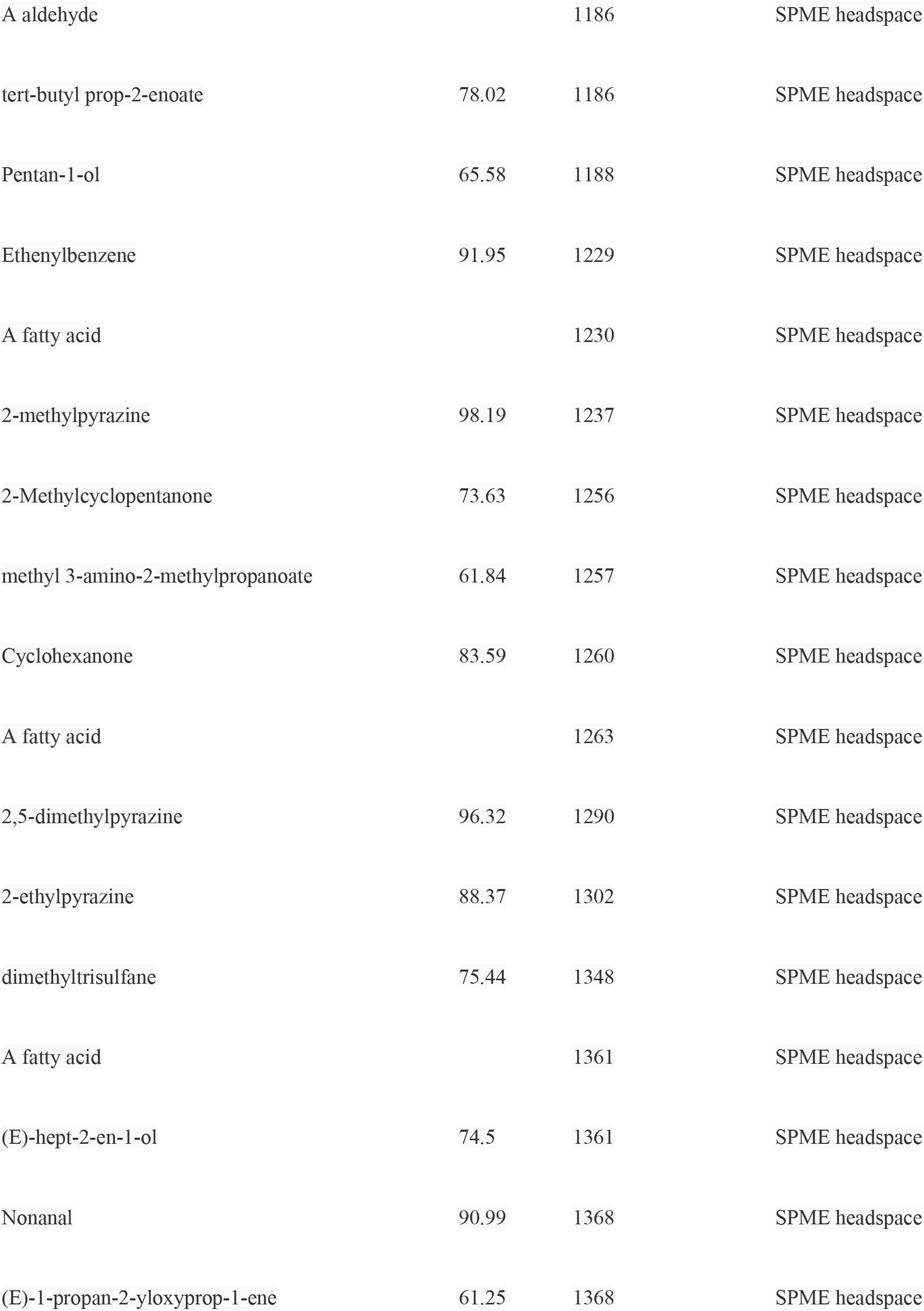
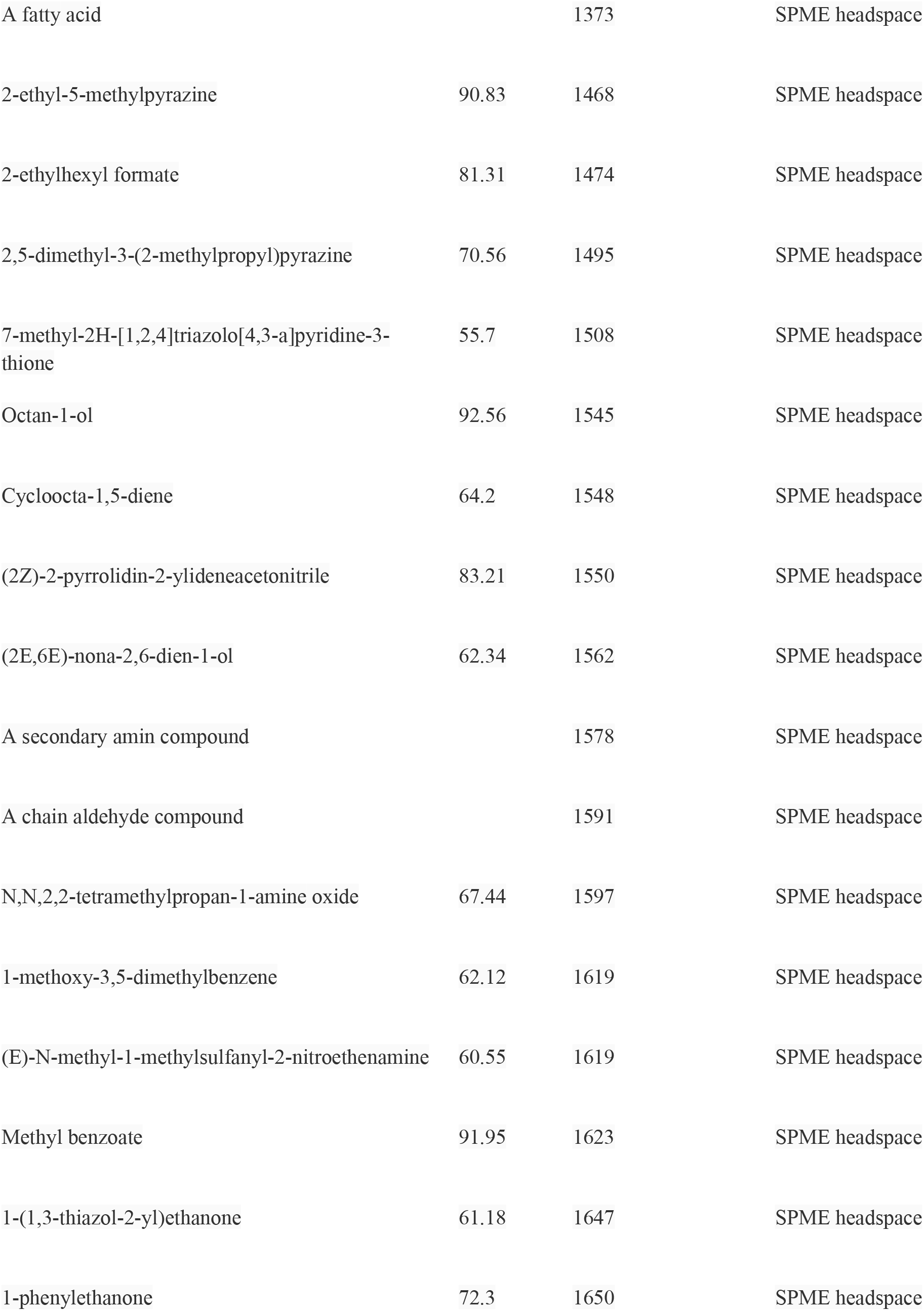
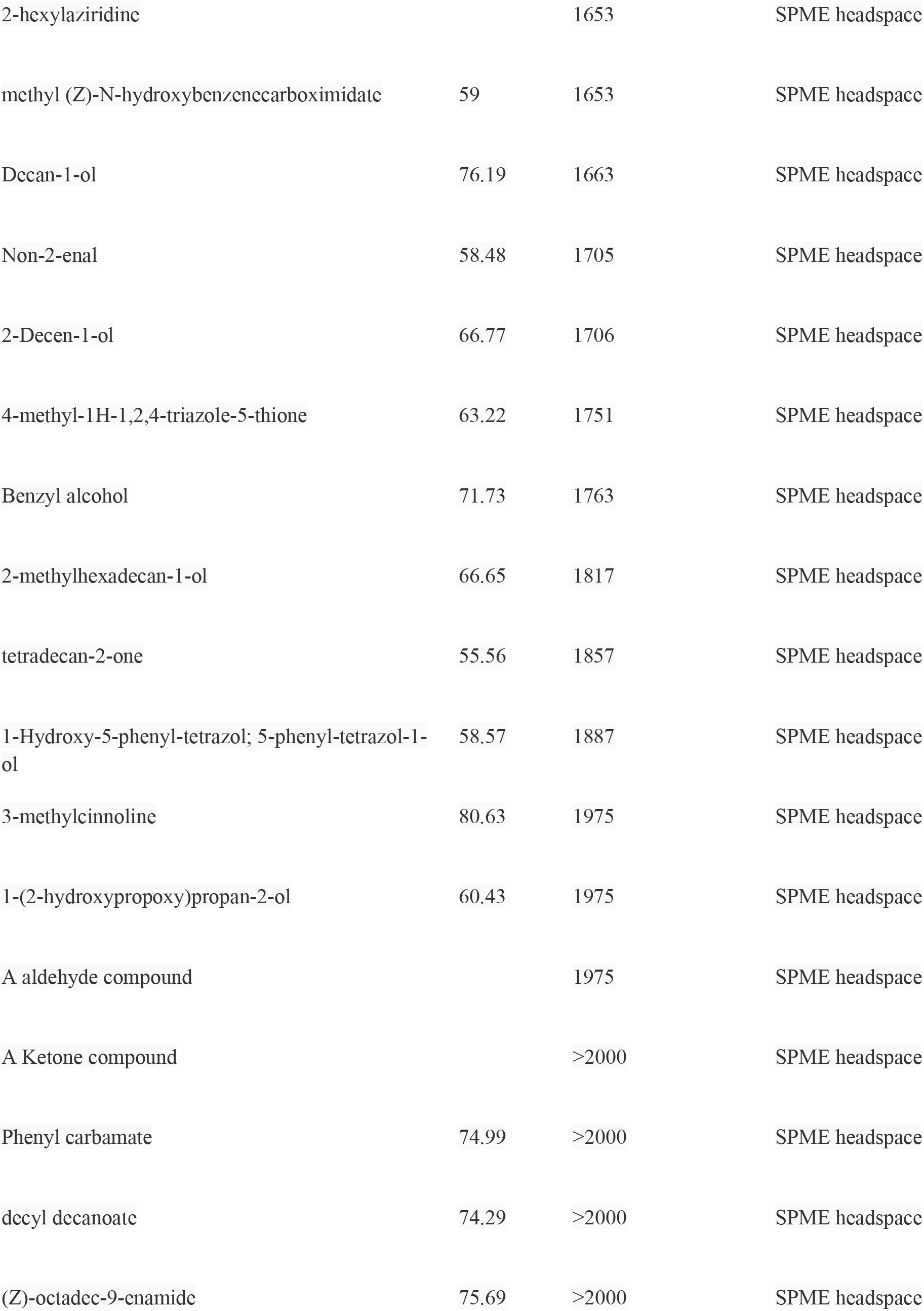
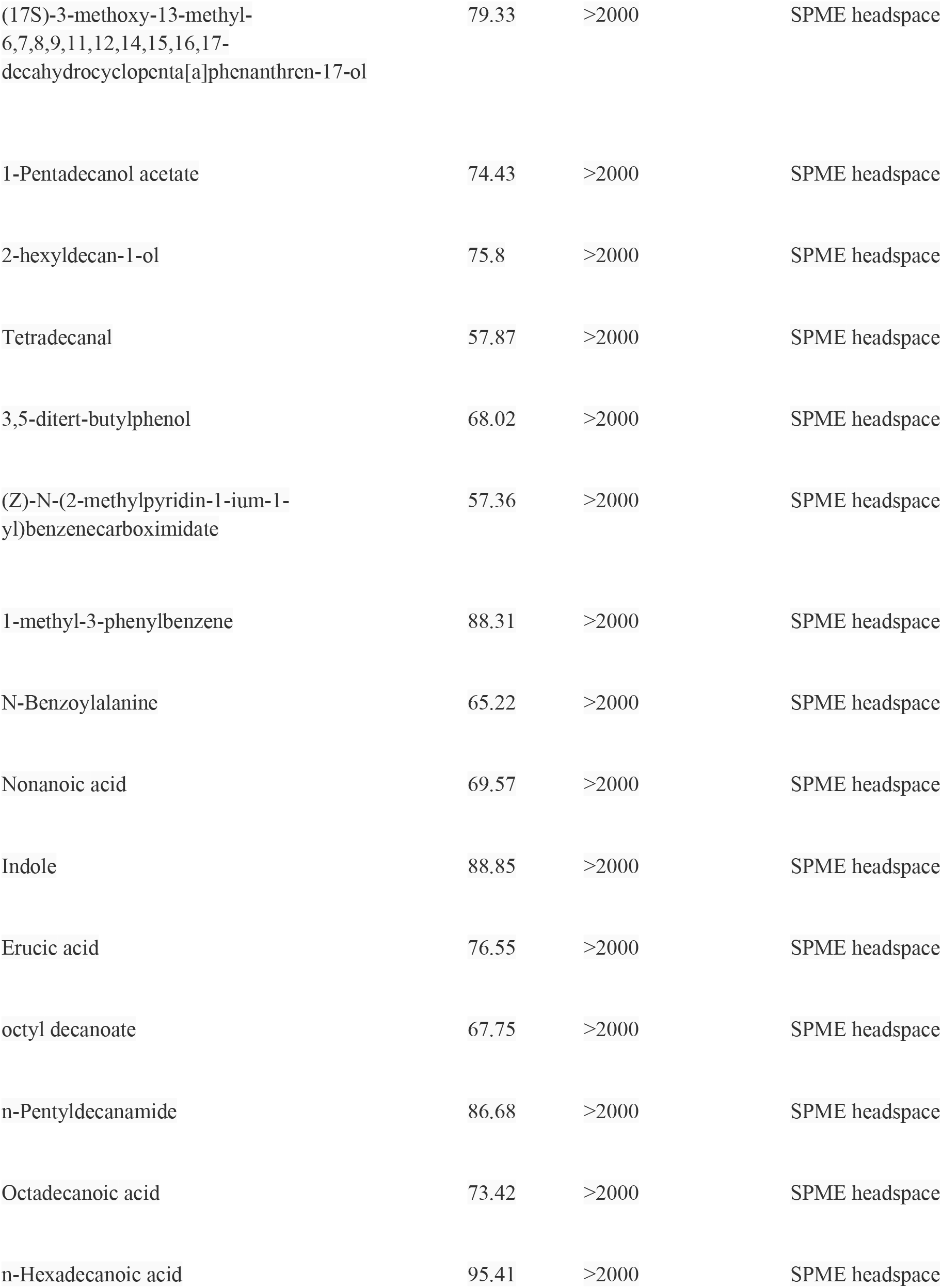
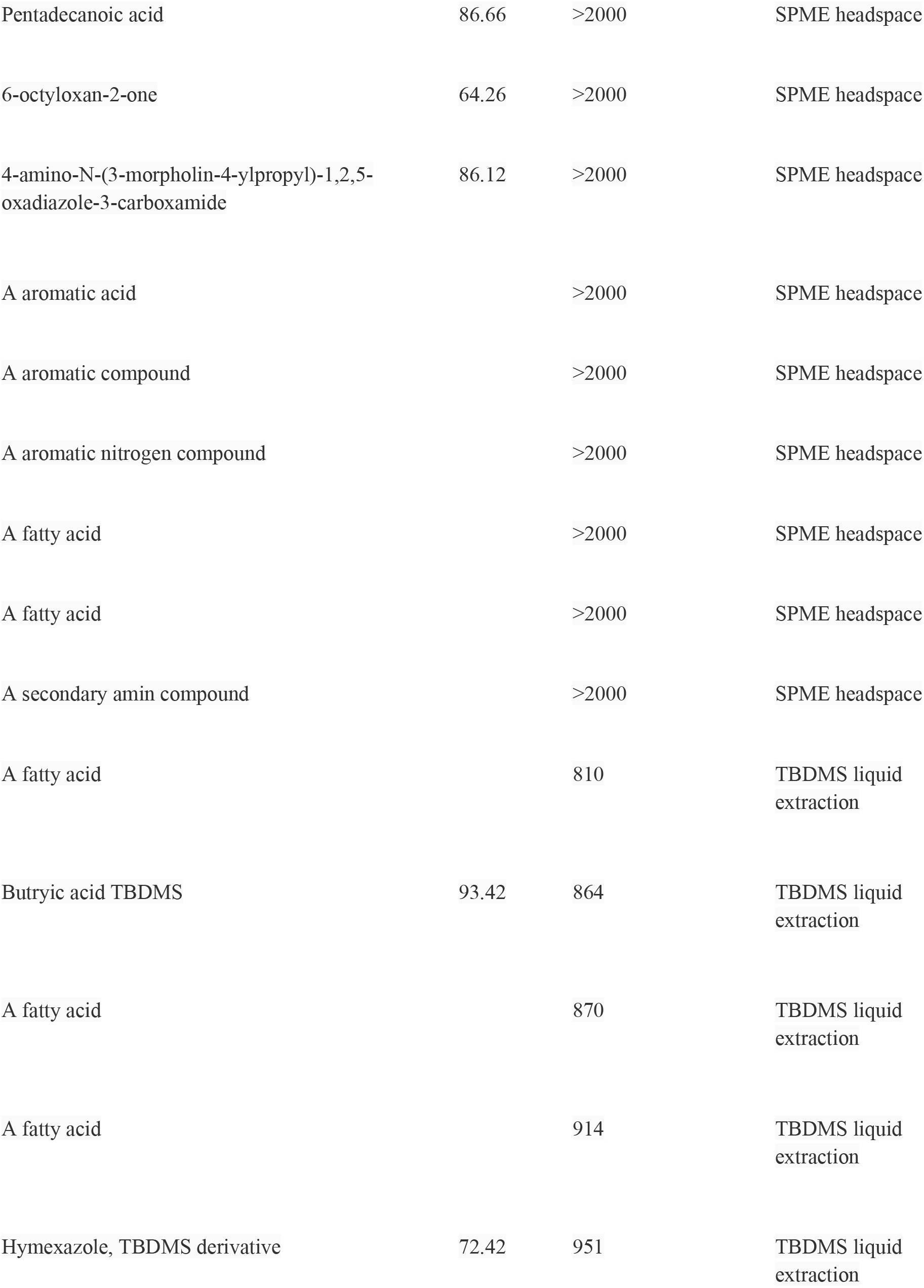
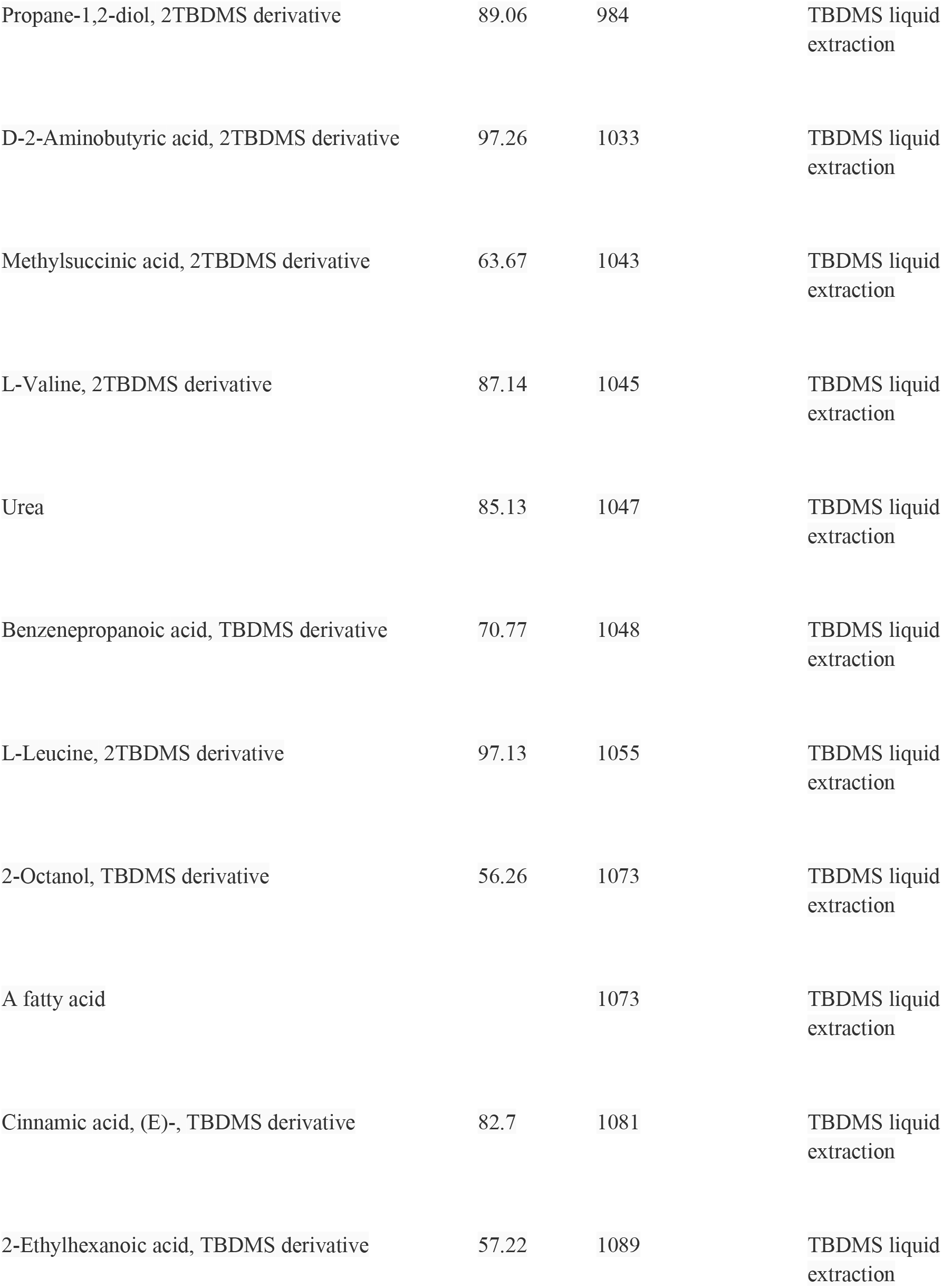
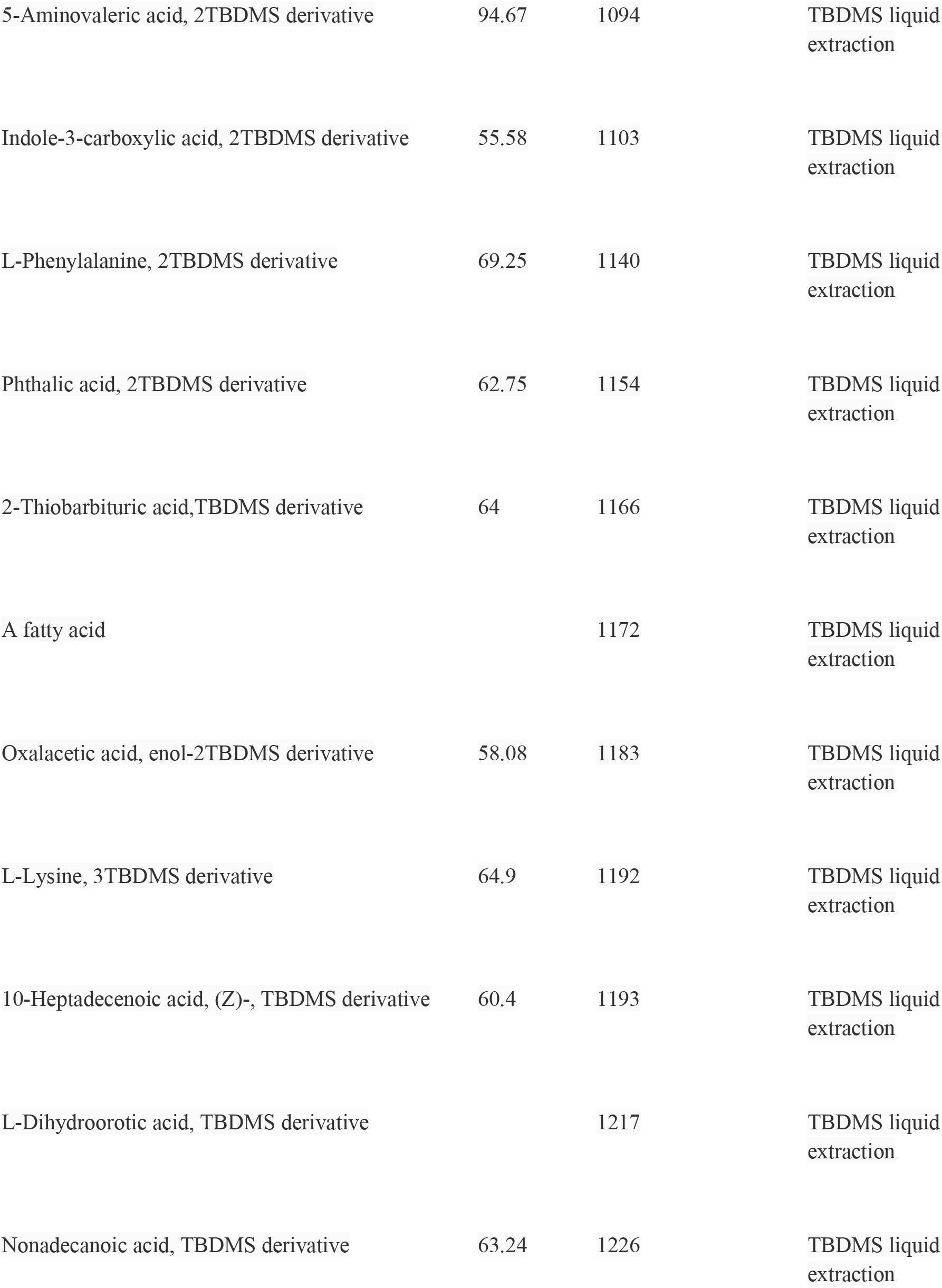
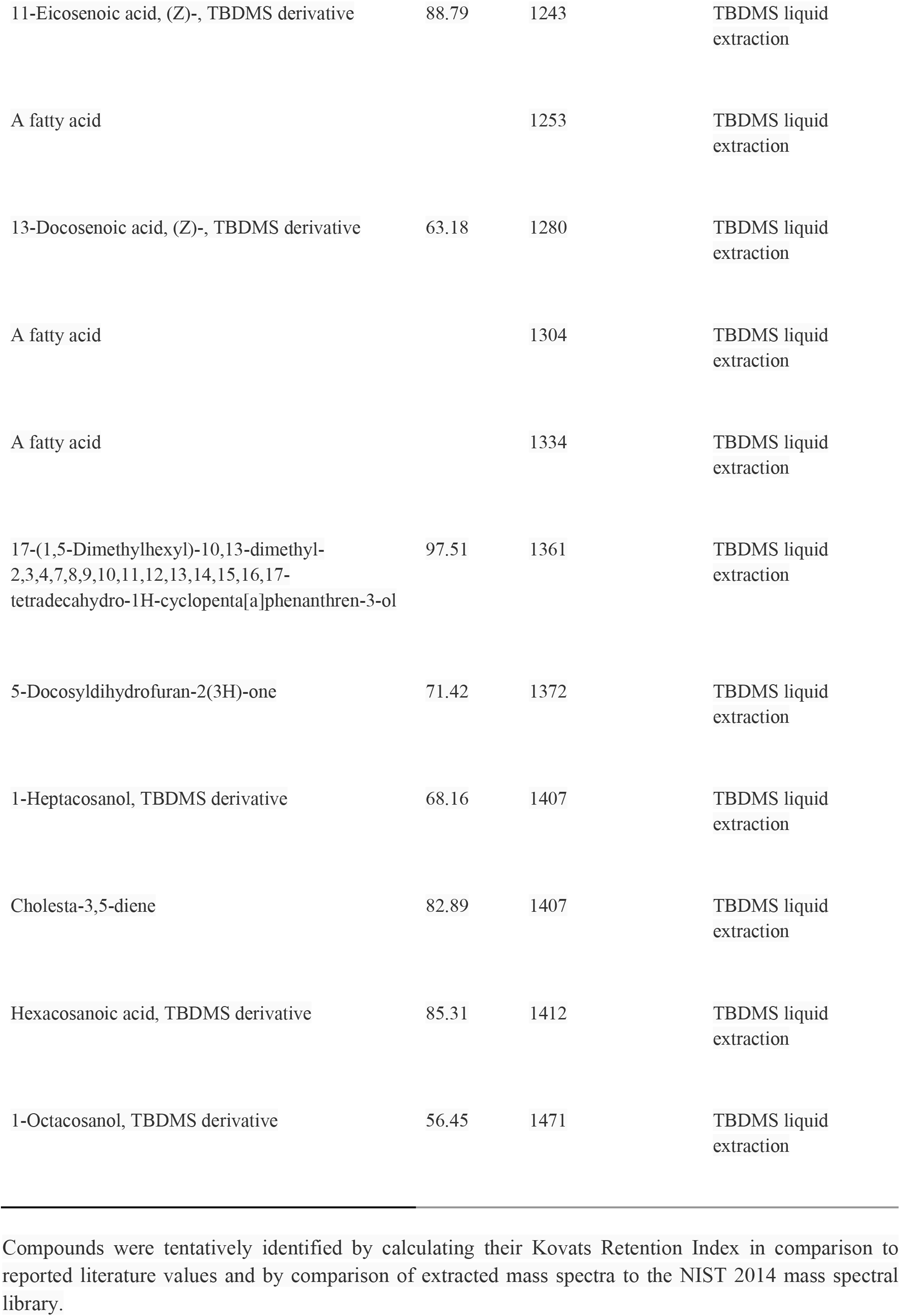
Tentatively identified compounds in the anal gland secretion of headspace solid phase microextraction (SPME) and liquid extraction tert-butyldimethylsilyl (TBDMS) derivative of domestic cat.

